# A Multisite and Multimodal Comparative Study of Cerebellar Connectome Between Multiple Sclerosis and Neuromyelitis Optica Spectrum Disorders

**DOI:** 10.1101/2022.01.16.476492

**Authors:** Yuping Yang, Junle Li, Zhen Li, Yaou Liu, Jinhui Wang

## Abstract

The cerebellum has been increasingly recognized to play key roles in the pathology of multiple sclerosis (MS) and spectrum disorders (NMOSD), two main demyelinating diseases with similar clinical presentations. Despite accumulating evidence from neuroimaging research for cerebellar volumetric alterations in the diseases, however, there have been no network-based studies examining convergent and divergent alterations in cerebellar connectome between MS and NMOSD. This multisite and multimodal study examined common and specific alterations in within-cerebellar coordination and cerebello-cerebral communication between MS and NMOSD by retrospectively collecting structural and resting-state functional MRI data from 208 MS patients, 200 NMOSD patients and 228 healthy controls (HCs) in seven sites in China. Morphological brain networks were constructed by estimating interregional similarity in cortical thickness and functional brain networks were formed by calculating interregional temporal synchronization in functional signals. After identifying cerebellar modular architecture and based on prior cerebral cytoarchitectonic classification and functional partition, within-cerebellar and cerebello-cerebral morphological and functional connectivity were compared among the MS, NMOSD and HC groups. Five modules were identified within the cerebellum including Primary Motor A (PMA), Primary Motor B (PMB), Primary Non-Motor (PNM), Secondary Motor (SM) and Secondary Non-Motor (SNM) modules. Compared with the HCs, the MS and NMOSD patients exhibited both increases and decreases in within-cerebellar morphological connectivity that were mainly involved in the PMA, PMB and SNM. Particularly, the two patient groups showed a common altered pattern characterized by decreases between the PMA and SNM, both of which were more densely connected with the PMB. For cerebello-cerebral morphological connectivity, widespread reductions were found in both patient groups for the SM and SNM with almost all cerebral cytoarchitectonic classes and functional systems while increases were observed only in the NMOSD patients for the PMB with cerebral areas involving motor and sensory domains. With regard to cerebellar functional connectivity, fewer alterations were observed in the patients that were all characterized by reductions and were mainly involved in cerebello-cerebral interactions between cerebellar motor modules and cerebral association cortex and high-order networks, particularly in the NMOSD patients. Cerebellar connectivity-based classification achieved around 60% accuracies to distinguish the three groups to each other with morphological connectivity as predominant features for differentiating the patients from controls while functional connectivity for discriminating the two diseases. Altogether, this study characterizes common and specific circuit dysfunctions of the cerebellum between MS and NMOSD, which provide novel insights into shared and unique pathophysiologic mechanisms underlying the two diseases.

## Introduction

While representing only about 10% of the size of the whole brain, the cerebellum comprises much more neurons than the cerebrum (Andersen, Korbo, and Pakkenberg 1992; Herculano-Houzel n.d.; Pelvig et al. 2008) and contains almost 80% of the surface area of the cerebral cortex (Sereno et al. 2020). The cerebellum has long been considered to devote exclusively to motor control, the last decade has however indicated that it is also engaged in various higher-level cognitive processes because of its complex internal structure and tight interactions with motor and non-motor regions of the cerebral cortex (Schmahmann et al. 2019; C. J. Stoodley and Schmahmann 2010). Moreover, cerebellar abnormalities are continuously demonstrated to be associated with a variety of motor or non-motor dysfunctions, such as gait ataxia, coordination deficiency, attention deficits, working memory decline, visual-spatial function damage and information processing speed droop (Hoppenbrouwers et al. 2008; Schmahmann et al. 2019). Thus, the cerebellum has attracted considerable research interests in recent years, particularly under different pathological conditions.

Multiple Sclerosis (MS) is an inflammatory and neurodegenerative disease affecting the central nervous system such as the optic nerve, spinal cord, and the cerebellum. Previous neuroimaging studies have shown that patients with MS exhibit widespread alterations in cerebellar structure and function including reduced grey matter volume (Anderson et al. 2009; Calabrese et al. 2010; Grothe et al. 2017; Ramasamy et al. 2009; Weier et al. 2012), decreased cerebellar regional homogeneity (Dogonowski et al., 2014) and altered cerebello-cerebral connectivity (Cerasa et al. 2012; Cocozza et al. 2018; Savini et al. 2019; Schoonheim et al. 2021; Tona et al. 2018). Neuromyelitis Optica spectrum disorders (NMOSD) is another inflammatory central nervous system disease that primarily affects the optic nerves and spinal cord with the IgG-antibody against the aquaporin-4 (AQP4) receptor as a specific autoantibody marker (Chang and Chang 2020; Lennon et al. 2004). Since the cerebellum has high AQP4 expression (Nico et al. 2002) and dense connections with the optic nerves and spinal cord, it has been increasingly investigated in NMOSD and is reported to show declined grey matter volume (Sun et al., 2019), disrupted microstructural integrity (Liu et al., 2018), impaired functional connectivity (Han et al., 2020; Liu et al., 2018) and abnormal spontaneous neural activity power (Liu et al., 2020) in patients. These findings collectively indicate that the cerebellum is a common, vulnerable structure to MS and NMOSD, which has important implications for understanding the physiopathology and improving the evaluation of clinical disability of the two diseases.

Despite the abovementioned advances, however, it is largely unknown whether differential patterns of structural and functional alterations occur in the cerebellum between MS and NMOSD. Currently, there are only a few studies that directly compare MS and NMOSD with respect to cerebellar alterations. These studies exclusively focused on cerebellar volume and consistently reported no significant differences between MS and NMOSD (Duan et al. 2021; Lee et al. 2018; Schneider et al. 2017; Weier et al. 2015). To date, there have been no network-based studies examining whether and how cerebellar structural and functional connectivity are differentially involved in MS and NMOSD and whether cerebellar connectivity have the potential to distinguish the two diseases.

In this study, we examined convergent and divergent alterations in cerebellum connectome by collecting structural and resting-state functional MRI data from a large cohort of participants in 7 sites in China. First, a multiplex network model was used to integrate morphological and functional connectivity for module detection within the cerebellum via a multilayer community detection algorithm. Then, cerebellar module-based morphological and functional connectivity were characterized within the cerebellum and between the cerebellum and cerebral cytoarchitectonic classes and functional systems. Finally, convergent and divergent alterations in cerebellar connectivity and their clinical and behavioral relevance as well as diagnostic power were examined.

## Materials and Methods

### Participants

This study included a total of 256 patients with MS, 270 patients with NMOSD and 281 health controls (HCs) from 7 different sites in China including Beijing Tiantan Hospital, Xuanwu Hospital, Tianjin Medical University General Hospital, Huashan Hospital, the First Affiliated Hospital of Nanchang University, China-Japan Union Hospital of Jilin University, and the First Affiliated Hospital of Chongqing Medical University. All participants were selected according to the following inclusion criteria: (1) the MS patients was diagnosed according to 2010 McDonald criteria (Polman et al. 2011) or 2017 revision of McDonald criteria (Thompson et al. 2018) and the NMOSD patients satisfied the recently 2015 NMOSD criteria (Wingerchuk et al. 2015); (2) without other neurological or psychiatric disorder; (3) the patients were either in relapsing phase (less than 4 weeks from the last relapse) or remitting phase (more than 4 weeks from the last relapse); (4) right-handed. We excluded 171 participants due to (1) image artifacts; (2) incomplete image or clinical data; and (3) excess head motion during the R-fMRI scan (see below for details). Finally, a total of 636 participants (208 MS patients, 200 NMOSD patients and 228 HCs) were included in the final analysis. This study was approved by the institutional review board of corresponding hospitals, and written informed consent was obtained from each participant.

### Clinical and neuropsychological assessment

Clinical variables included disease duration and Expanded Disability Status Scale (EDSS). For neuropsychological assessment, the following tests were used: the California Verbal Learning Test-Second Edition (CVLT; accessing the auditory or verbal episodic memory), the Brief Visuospatial Memory Test-Revised (BVMT, accessing the visual or spatial episodic memory) and the Paced Auditory Serial Addition Task (PASAT; accessing auditory processing speed, attention and working memory). Of note, participants in only 3 sites (Beijing Tiantan Hospital, Xuanwu Hospital, and Tianjin Medical University General Hospital) completed the neuropsychological tests.

### Multimodal MR image acquisition

Multimodal images were acquired in this study, including T2 FLAIR (fluid-attenuated inversion recovery) image, high-resolution 3D T1-weighted imaging and R-fMRI. The imaging parameters are summarized in Table S1.

### WM lesion loads and structural image filling

For each patient, a WM lesion mask was manually delineated using the 3D-slicer software (https://www.slicer.org) if WM hyperintensity was evident on his or her T2 FLAIR image. Individual WM lesion loads were calculated as the total volumes within the masks. Based on the masks, a SLF algorithm was used to refill WM lesions on individual T1 images (http://atc.udg.edu/salem/slfToolbox/software.html).

### Structural MRI processing

#### Cerebellar cortical thickness

Cortical thickness estimation of the cerebellum was accomplished by a patch-based multi-atlas segmentation tool called CERES (Romero et al. 2017) *-* the winner of the Medical Image Computing and Computer-Assisted Intervention cerebellum segmentation challenge (Carass et al. 2018) *-* on the web-based platform volBrain (http://volbrain.upv.es; (Manjón and Coupé 2016). Briefly, a spatially adaptive non-local means filter was first applied to reduce noise in the structural images. Then, the N4 bias field correction (Tustison et al. 2010) was performed to correct intensity inhomogeneity. The corrected images were further linearly registered (affine transform) to the standard Montreal Neurological Institute (MNI) space using the Advanced Normalization Tools, followed by the N4 bias field correction again. Subsequent analyses were limited to the cerebellar areas to reduce the computational burden. To achieve a better cerebellum anatomic matching between the corrected images and the library of manually labeled templates, the non-linear transformations were estimated for the corrected images and templates to the MNI152 atlas. By concatenating the forward non-linear transformation to the MNI152 atlas of each library case and the inverse non-linear transformation of each corrected image, a subject-specific library was obtained. Finally, a local intensity normalization based on regions of interest (ROIs) derived from majority voting segmentation of the subject-specific library was applied to ensure the same intensity of cerebellar tissues across participants.

#### Cerebral cortical thickness

Cortical thickness estimation of cerebral cortex was performed by the CAT12 toolbox (http://dbm.neuro.uni-jena.de/cat12/) based on the SPM12 package (https://www.fil.ion.ucl.ac.uk/spm/software/spm12/). As an efficient and reliable alternative to FreeSurfer, the CAT12 toolbox provides a volume-based approach for cortical thickness estimation without extensive reconstruction of cortical surface. The CAT12 started with an initial segmentation of individual structural images into gray matter (GM), white matter (WM) and cerebrospinal fluid (CSF) based on an adaptive Maximum A Posterior technique (Rajapakse, Giedd, and Rapoport 1997). Then, a projection-based mothed (Dahnke, Yotter, and Gaser 2013) was performed to estimate cortical thickness which could handle the partial volume information, sulcal blurring and sulcal asymmetries. Subsequently, a central cortical surface was created and reparametrized into a common coordinate system via spherical mapping (Yotter, Thompson, and Gaser 2011). Finally, individual cortical thickness maps were resampled into the common fsaverage template and smoothed using a Gaussian kernel with 15-mm full width at half maximum.

### Functional MRI processing

The R-fMRI data were processed with the GRETNA toolbox (J. Wang et al. 2015) based on the SPM12 package (http://www.fil.ion.ucl.ac.uk/spm/software/spm12/). First, following removal of the first 5 volumes for magnetic saturation, inter-volume head motion was corrected via rigid transformation. After converting rotational displacements from degrees to millimeters on the surface of a sphere of radius 50 mm (Power et al. 2012), participants with excessive motion were excluded in terms of the criteria of > 3 mm translation or > 0.5 mm mean frame-wise displacement. For the remaining participants, there were no significant differences in the maximum [HCs = 0.675 (IQR = 0.620), MS = 0.688 (IQR = 0.600), and NMOSD = 0.641 (IQR = 0.552); *p* = 0.287, permutation test] and mean frame-wise [HCs = 0.150 (IQR = 0.102), MS = 0.137 (IQR = 0.099), and NMOSD = 0.139 (IQR = 0.116); *p* = 0.529, permutation test] displacement of head motion among three groups. Then, the corrected functional images underwent band-pass filtering (0.01 - 0.08 Hz) and nuisance regression [24-parameter head motion profiles (Friston et al. 1996), white matter signals, cerebrospinal fluid signals and global signals] in a single linear model to avoid reintroducing artifacts (Lindquist et al. 2019). The white matter and cerebrospinal fluid signals were calculated within subject-specific masks derived from tissue segmentation of individual structural images (threshold = 0.9), which were co-registered to the corresponding mean volume of the corrected functional images. Finally, the functional images were normalized into standard MNI space by applying deformation fields derived from the tissue segmentation of individual structural images, and spatially smoothed by a Gaussian kernel (full width at half maximum = 6 mm).

### Construction of within-cerebellar networks

#### Cerebellar parcellation

In the CERES, the cerebellum was segmented using a multi-atlas segmentation technique called non-local patch-based label fusion (Coupé et al. 2011). Specifically, an Optimized PatchMatch algorithm (Giraud et al. 2016; Ta et al. 2014) was performed to speed up the patch matching process and an adaptive multi-scale approach was used for better accuracy. To avoid slightly irregular edges in the resultant label maps, a convolution with a smoothing kernel of 5 × 5 × 5 voxels was further applied to the label maps. Finally, the cerebellum of each participant was divided into 24 regions including the lobule I_II, lobule III, lobule IV, lobule V, lobule VI, Crus I, Crus II, lobule VIIb, lobule VIIIa, lobule VIIIb, lobule IX and lobule X in each hemisphere (Figure 1).

**Figure 1.**
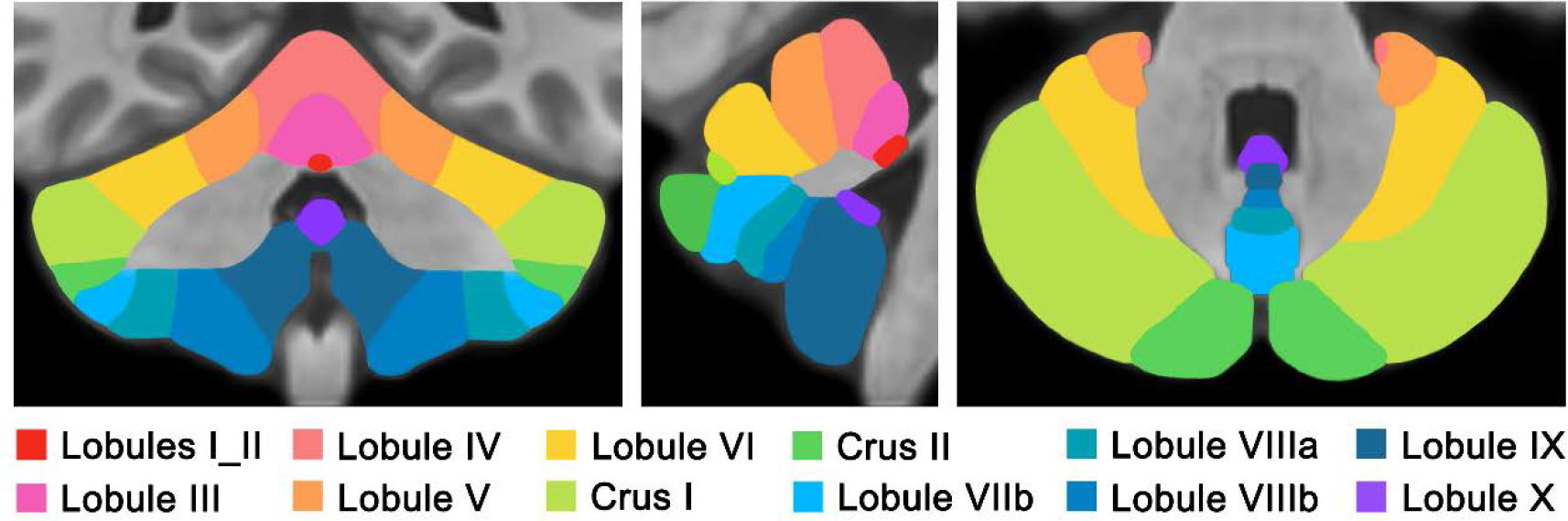
Illustrative representation of cerebellar lobules. According to the CERES partition, the cerebellum was segmented into 12 lobules in each hemisphere.

#### Morphological connectivity estimation

For a given pair of regions within the cerebellum, morphological connectivity was estimated by calculating Jensen-Shannon divergence based similarity in the distribution of intraregional cortical thickness. Details on the calculation of morphological connectivity are described elsewhere (Li et al. 2021; H. Wang et al. 2016).

#### Functional connectivity estimation

For a given pair of regions within the cerebellum, functional connectivity was estimated by calculating Pearson correlation in intraregional mean time series.

### Construction of cerebello-cerebral networks

#### Cerebral parcellation

The cerebrum was divided into 400 ROIs using a functionally defined atlas, which was derived from R-fMRI data from 1489 subjects (Schaefer et al. 2018).

#### Morphological connectivity estimation

For each pair of regions between the cerebellum and cerebrum, morphological connectivity was estimated using the same methods as for the cerebellar morphological networks.

#### Functional connectivity estimation

For each pair of regions between the cerebellum and cerebrum, functional connectivity was estimated using the same methods as for the cerebellar functional networks.

### Removal of multicenter effects

For multicenter studies, a crucial step is to remove site effects to avoid spurious findings (Han et al. 2006). In this study, we utilized a harmonization approach called Combat (J. P. Fortin et al. 2018) to moderate site effects. The Combat harmonization is demonstrated to successfully remove inter-site technical variability while preserving inter-site biological variability in image-based measurements. Specifically, the Combat model can be written as:

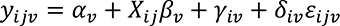

where *y_*i*jv_* represents connectivity strength of edge *v* (or cortical thickness of region *v*) for subject *j* in site *i*, *α*_*v*_ is the average connectivity strength (or cortical thickness) for edge (or region) *v*, *X* is a design matrix for the covariates of interest (e.g., age, sex and group), *β*_*v*_ is a vector of regression coefficients corresponding to covariates in *X*, and *ε_*i*j*v*_* is the residual term that is assumed to follow a normal distribution with zero mean. The terms *γ*_*iv*_ and *δ*_*iv*_ represent the additive and multiplicative site effects of site *i* on edge (or region) *v*, respectively, and are estimated by conditional posterior means as described in (J.-P. Fortin et al. 2017; Johnson, Li, and Rabinovic 2007). The final ComBat-harmonized connectivity strength (or cortical thickness) for edge (or region) *v* is calculated as:

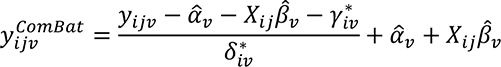

where 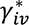 and 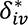 are the empirical Bayes estimates of *γ_*iv*_* and *δ_*iv*_*, respectively.

### Cerebellar module detection

Cerebellar modular architecture was identified by applying a multilayer community detection algorithm to a group-level multiplex network, which integrated both morphological and functional connectivity within the cerebellum for the HCs. First, a group-level mean morphological network and functional network were separately obtained for the HCs. As two layers, these two networks were then interconnected via adding edges to link each node in one layer with replica of the node in the other layer, therefore forming a multiplex network. Thereafter, a nonparametric method of locally adaptive network sparsification (Foti, Hughes, and Rockmore 2011) was utilized such that only those locally significant connections that could not be explained by random variation were retained in the multiplex network. Module detection is to find a specific partition of network nodes that yields the largest modularity, *Q*, which is defined for a multilayer network as (Mucha, Richardson, Macon, Porter, & Onnela, 2010):

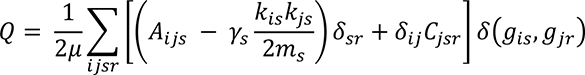

where *i* and *j* represent nodes, *s* and *r* represent layers, *A_ijs_* denotes intra-layer connectivity strength between node *i* and node *j* in layer *s*, *C_jsr_* denotes inter-layer connectivity strength of node *j* between layer *s* and layer *r*, *k*_*is*_ = ∑*_j_ A*_*ijs*_ denotes total intra-layer connectivity strength of node *i* in layer *s*, *m_s_* is total connectivity strength of all edges in layer *s*, *γ_s_* is module resolution in layer *s* (the larger *γ*, the more small-size communities), *g_is_*/*g_jr_* represents module assignment of node *i*/*j* in layer *s*/*r*, 2*μ* is equal to ∑*_jr_*(*k_js_* + *c_js_*) with *c_js_* = ∑*_r_ C_jsr_* indicating total inter-layer connectivity strength node *j* from layer *s* to other layers, and *δ* is a binary notation (1 if the same node for *δ_*i*j_*, the same layer for *δ*_*sr*_ and the same community assignment *δ*(*g_*i*s_*, *g_jr_*) and 0 otherwise). In this study, the module detection was performed using the Louvain algorithm (Blondel et al. 2008).

During the processes mentioned above, there were two parameters that may affect final module identification: the inter-layer connectivity strength, *ω*, and the module resolution, *γ*. To identify stable cerebellar modular architecture, we calculated the modularity, *Q* across a large range in the 2D parameter space of (*ω*, *γ*): *ω* = [0.01 - 1] and *γ* = [0.01 - 3] both with an increment of 0.01. Specifically, at each (*ω*, *γ*) in the 2D parameter space, we performed module detection 1,000 times and obtained a consensus matrix in which the elements indicated the proportions of each pair of nodes that were assigned to the same module over the 1,000 repetitions (Bassett et al. 2013). If there existed any pair of nodes that were not consistently assigned to the same module, the module detection was further conducted on the consensus matrix (1,000 times) to derive a new consensus matrix. Such procedures were iterated until a consensus partition was obtained. That is, the final consensus matrix included only 0 and 1. After determining cerebellar partition at each (*ω*, *γ*), we employed the variation of information (Meilă 2003) to evaluate stability of the cerebellar partition over different (*ω*, *γ*). More specifically, we computed the mean variation of information of cerebellar partition between a given (*ω*_*i*_, *γ*_*i*_) and its eight contiguous neighbors 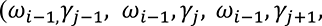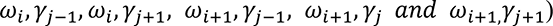. The smaller the mean variation of information was, the more stable the cerebellar partition was when subtle (*ω*, *γ*) fluctuations occurred. Finally, we searched the widest continuous (*ω*, *γ*) range in which the mean variation of information was equal to the minimum, and the corresponding cerebellar partition was regarded as canonical cerebellar modular architecture.

### Cortical cytoarchitectonic classification and functional partition

The cerebrum can be divided into several canonical modules based on different approaches. In this study, two partitions *-* cytoarchitectonic classification and functional partition *-* were used to divide the cerebrum since a recent study showed that structural and functional connectivity followed cytoarchitectonic and functional hierarchies in the cerebrum (Vázquez-Rodríguez et al. 2019). 1) Cytoarchitectonic classification of the cerebrum (cerebrum^C^ hereafter). The cerebral cortex displays substantial variation in cytoarchitecture, according to which the cerebrum is categorized into seven classes (Seidlitz et al. 2018; Vértes et al. 2016): primary motor cortex (PM), association cortex (AC1), association cortex (AC2), primary/secondary sensory (PSS), primary sensory cortex (PS), limbic regions (LB) and insular cortex (IC). We manually assigned each of the 400 cerebral regions to one of the seven cytoarchitectonic classes. 2) Functional partition of the cerebrum (cerebrum^F^ hereafter). In terms of intrinsic functional connectivity patterns, the second partition divides the cerebrum into seven functional systems (Thomas Yeo et al. 2011): visual network (VN), somatomotor network (SMN), dorsal attention network (DAN), ventral attention network (VAN), limbic network (LN), fronto-parietal network (FPN) and default mode network (DMN). Each of the 400 cerebral regions belonged to one of the seven functional systems.

### Construction of cerebellar module**-**based networks within the cerebellum and between the cerebellum and cerebrum

Based on the methods mentioned above, we constructed 4 networks for each participant at the region level: one 24 × 24 within-cerebellar morphological network, one 24 × 24 within-cerebellar functional network, one 24 × 400 cerebello-cerebral morphological network and one 24 × 400 cerebello-cerebral functional network. Based on the cerebellar modular architecture (5 modules; see Results for details) and cerebrum^C^ and cerebrum^F^, the 4 region-level networks were transformed to 6 cerebellar module-based networks: one 5 × 5 within-cerebellar morphological network, one 5 × 5 within-cerebellar functional network, one 5 × 7 cerebello-cerebral^C^ morphological network, one 5 × 7 cerebello-cerebral^F^ morphological network, one 5 × 7 cerebello-cerebral^C^ functional network and one 5 × 7 cerebello-cerebral^F^ functional network. This was achieved by averaging connectivity weights of all connections linking any pair of cerebellar modules or linking one cerebellar module and one cerebral cytoarchitectonic class/functional system.

### Statistical analysis

#### Group effects

Chi-squared tests were used to compare dichotomous variables including sex (male vs. female), disease state (acute vs. chronic) and WM lesion (presence vs. absence). For continuous variables (demographic: age; clinical: disease duration, EDSS and WM lesion volume; neuropsychological: CVLT, BVMT and PASAT; imaging-based: cortical thickness, morphological connectivity and functional connectivity), non-parametric permutation tests were utilized after examining their normality (Lilliefors test). Age, sex and mean framewise displacement of head motion (if applicable) were treated as covariates in the analyses of neuropsychological variables and imaging-based measures. Briefly, for a given continuous variable, we initially calculated a statistic (T or F) through two-sample t test (for disease duration, EDSS and WM lesion volume), ANOVA (for age) or ANCOVA (for CVLT, BVMT, PASAT, cortical thickness, morphological connectivity and functional connectivity). To obtain an empirical distribution of the statistic, we randomly reshuffled the data and re-computed the statistic (10,000 times). Based on the empirical distribution, a *p* value was calculated as the proportion of permutations that generated the statistic equal to or greater or less than (for T) or equal to or greater than (for F) the real observation. Notably, the covariates were not reshuffled during the permutation process. To correct for multiple comparisons for imaging-based measures, the false discovery rate (FDR) procedure was used at the level of *q* < 0.05 (cortical thickness: 5 cerebellar modules; morphological/functional connectivity: 15 within-cerebellar and 35 cerebello-cerebral connections). For significant differences among the three groups, post hoc pairwise comparisons were further performed by permutation tests.

#### Relationship between cerebellar alterations and clinical/neuropsychological variables

For imaging-based metrics showing significant group effects, non-parametric Spearman partial correlation was used to evaluate their relationships with clinical and neuropsychological variables in the MS and/or NMOSD group in terms of patterns of post hoc comparisons. Age, sex and mean framewise displacement of head motion (if applicable) were regarded as covariates in the correlation analyses. Again, the FDR procedure was employed to correct for multiple comparisons.

### Classification analysis

To examine the potential of cerebellar connectivity in distinguishing the three groups from each other, we trained linear SVM classifiers with all within-cerebellar and cerebello-cerebral morphological and functional connectivity as initial features. Out-of-sample classification performance of the classifiers was evaluated using a 10-fold cross-validation procedure. Specifically, in each fold of classification between two groups, feature selection was first performed to exclude irrelevant features via non-parametric permutation tests on the training dataset (*p* < 0.05; 1,000 permutations). Based on the selected features, a linear SVM classifier was then trained and applied to the testing dataset. After the 10-fold cross-validation procedure, a classification accuracy was calculated. To obtain robust estimation of the classification accuracy, the above classification process was repeated 100 times and the resultant mean across all repeats was reported. Meanwhile, the consensus features that were consistently selected in at least 90% of all folds and repeats (10 × 100 = 1,000) were recorded together with their mean weights. Finally, to further evaluate whether the classifiers performed significantly better than random operations, an empirical null distribution of classification accuracy was obtained by reshuffling the group labels of participants (1,000 times) followed by 10-fold cross-validated classification. A *p* value was computed as the proportion of values in the empirical null distribution that were larger than the real observation. Notably, effects of age, sex, and mean frame-wise head motion (if applicable) were removed from all features via multiple linear regression before the classification procedure.

## Results

### Demographic, clinical, and neuropsychological evaluations

Table 1 summarizes demographic, clinical, and neuropsychological information of the participants. Significant group effects were found on age (*p* = 0.003) and sex (*p* < 0.001) separately due to an older age of the NMOSD patients than the MS patients (*p* < 0.001) and HCs (*p* = 0.020) and a higher female-to-male ratio of the NMOSD patients than the MS patients (*p* < 0.001) and HCs (*p* < 0.001) as well as of the MS patients than the HCs (*p* = 0.025). For clinical data, no significant differences were found in the acute-to-chronic phase ratio or disease duration between the two patient groups (*p* > 0.05). However, the NMOSD patients exhibited lower incidence of WM lesion (*p* < 0.001) and less lesion volume (*p* < 0.001) than the MS patients. In addition, the NMOSD patients showed significantly higher EDSS than the MS patients (*p* < 0.001). Regarding neuropsychological variables, both the patient groups performed significantly worse than the HCs in the CVLT and PASAT (both *p* < 0.001). In addition, the NMOSD patients showed lower BVMT than the HCs (*p* < 0.001). No differences were found in any neuropsychological variable between the two patient groups (*p* > 0.05).

**Table 1.** Demographic, clinical and neuropsychological variables.

|  | HCS<br>(n = 228) | MS<br>(n = 208) | NMO<br>(n = 200) | *p*-value |
| --- | --- | --- | --- | --- |
| Age (years) | 37.0 (21.0) | 36.0 (17.0) | 41.0 (21.5) | 0.003<sup>a,c</sup> |
| Sex (female/male) | 124/104 | 135/73 | 175/25 | <0.001<sup>a,b,c</sup> |
| Disease state (acute/chronic)* | - | 48/149 | 48/147 | 0.954 |
| Disease duration (months) | - | 19.0 (54.0) | 36.0 (52.0) | 0.054 |
| Lesion |  |  |  |  |
| N (%) | - | 208 (100%) | 110 (55%) | <0.001 |
| Volume (cm<sup>3</sup>) | - | 7.6 (16.0) | 1.2 (4.0) | <0.001 |
| EDSS | - | 2.0 (2.5) | 3.5 (3.0) | <0.001 |
| CVLT<sup>†</sup> | 52.0 (10.8) | 47.5 (15.0) | 47.0 (14.5) | <0.001<sup>b,c</sup> |
| PASAT<sup>†</sup> | 51.5 (14.0) | 40.0 (17.0) | 40.0 (18.0) | <0.001<sup>b,c</sup> |
| BVMT<sup>†</sup> | 28.0 (5.8) | 27.0 (11.8) | 22.0 (12.0) | 0.003<sup>c</sup> |

Data are represented as median (interquartile range) unless stated otherwise. HCs, healthy controls; MS, multiple sclerosis; NMO, neuromyelitis optica; EDSS, Expanded Disability Status Scale; CVLT, California Verbal Learning Test; PASAT, Paced Auditory Serial Addition Test; BVMT, Brief Visuospatial Memory Test.
\*Data are missing for 16 patients.
<sup>†</sup>Data are available only for a subset of participants from three hospitals (see Materials and Methods for details).
<sup>a</sup>Significant differences between the two patient groups.
<sup>b</sup>Significant differences between the MS patients and HCs.
<sup>c</sup>Significant differences between the NMO patients and HCs.

### Cerebellar modular architecture

In the searched 2D parameter space of inter-layer connectivity strength, *ω*, and resolution coefficient, *γ*, we identified a widest range (*ω* = 0.06 ∼ 0.50; *γ* = 1.39 ∼ 2.00) wherein cerebellar modular architecture maintained stable when the parameters fluctuated (Figure 2). Specifically, for each combination of the parameter pair in the identified range, the cerebellum was subdivided into five spatially contiguous and bilaterally symmetrical modules with regions resembling largely between morphological and functional networks (*Q* = 0.566; Figure 3). The modules had a good correspondence with the well-established cerebellar double-motor/triple-non-motor organization, and thus were termed Primary Motor A module (PMA, including bilateral Lobule I-II and Lobule III, with additional bilateral Lobule X for morphological networks), Primary Motor B module (PMB, including bilateral Lobule IV, Lobule V and Lobule VI), Primary Non-Motor module (PNM, including bilateral Crus I and Crus II), Secondary Motor module (SM, including bilateral Lobule VIIb, Lobule VIIIa and Lobule VIIIb) and Secondary Non-Motor module (SNM, including bilateral lobule IX, with additional bilateral Lobule X for functional networks).

**Figure 2.**
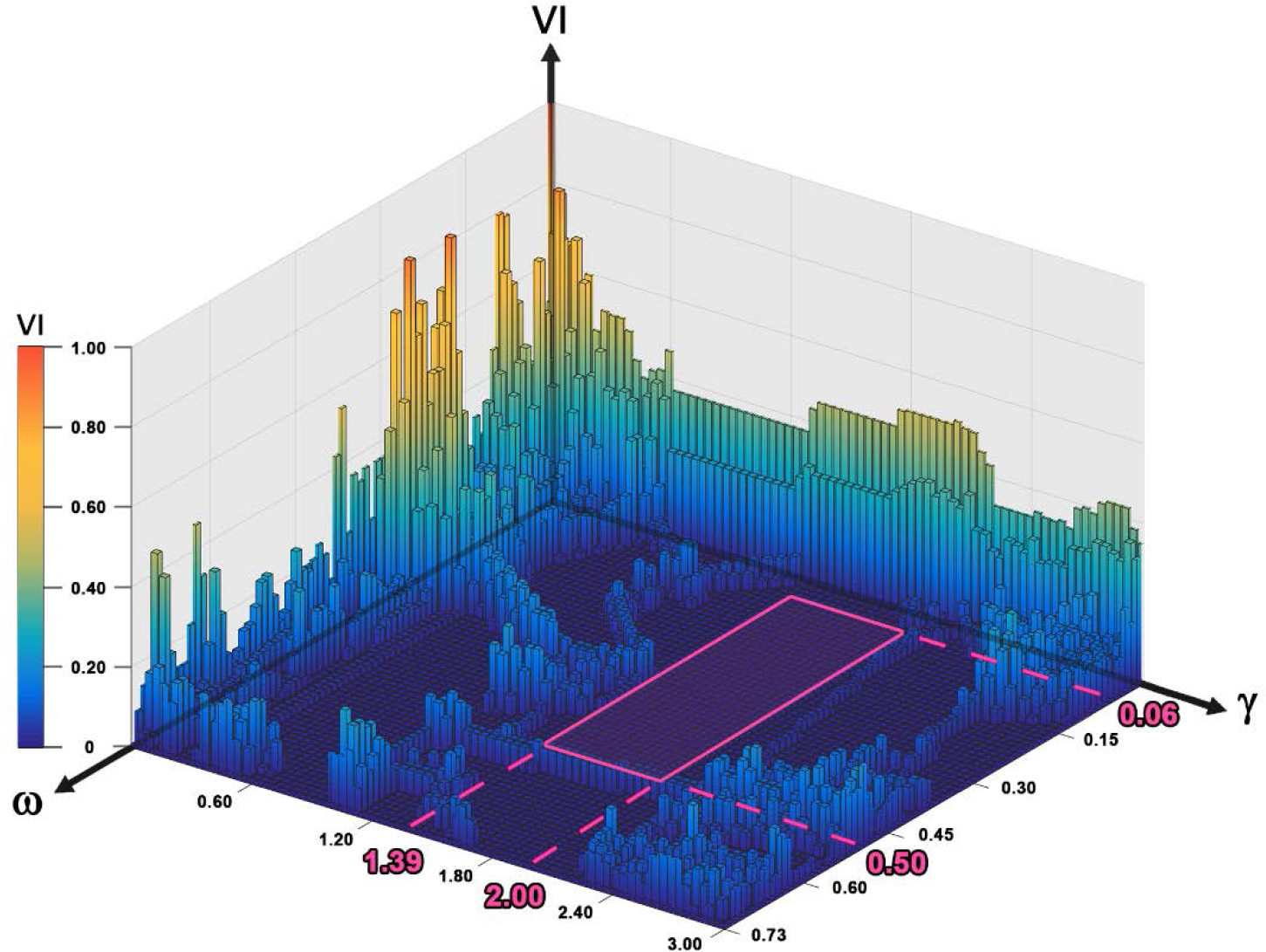
The stability of cerebellar modular architecture over different combinations of inter-layer connectivity strength (*ω*) and the module resolution (*γ*). The variation of information was used to evaluate the stability of cerebellar modular architecture. In the searched 2D parameter space (*ω* = [0.01 - 1]; *γ* = [0.01 - 3]), a widest range (*ω* = 0.06 ∼ 0.50; *γ* = 1.39 ∼ 2.00) was identified wherein cerebellar modular architecture maintained stable when the parameters fluctuated. VI, variation of information.

**Figure 3.**
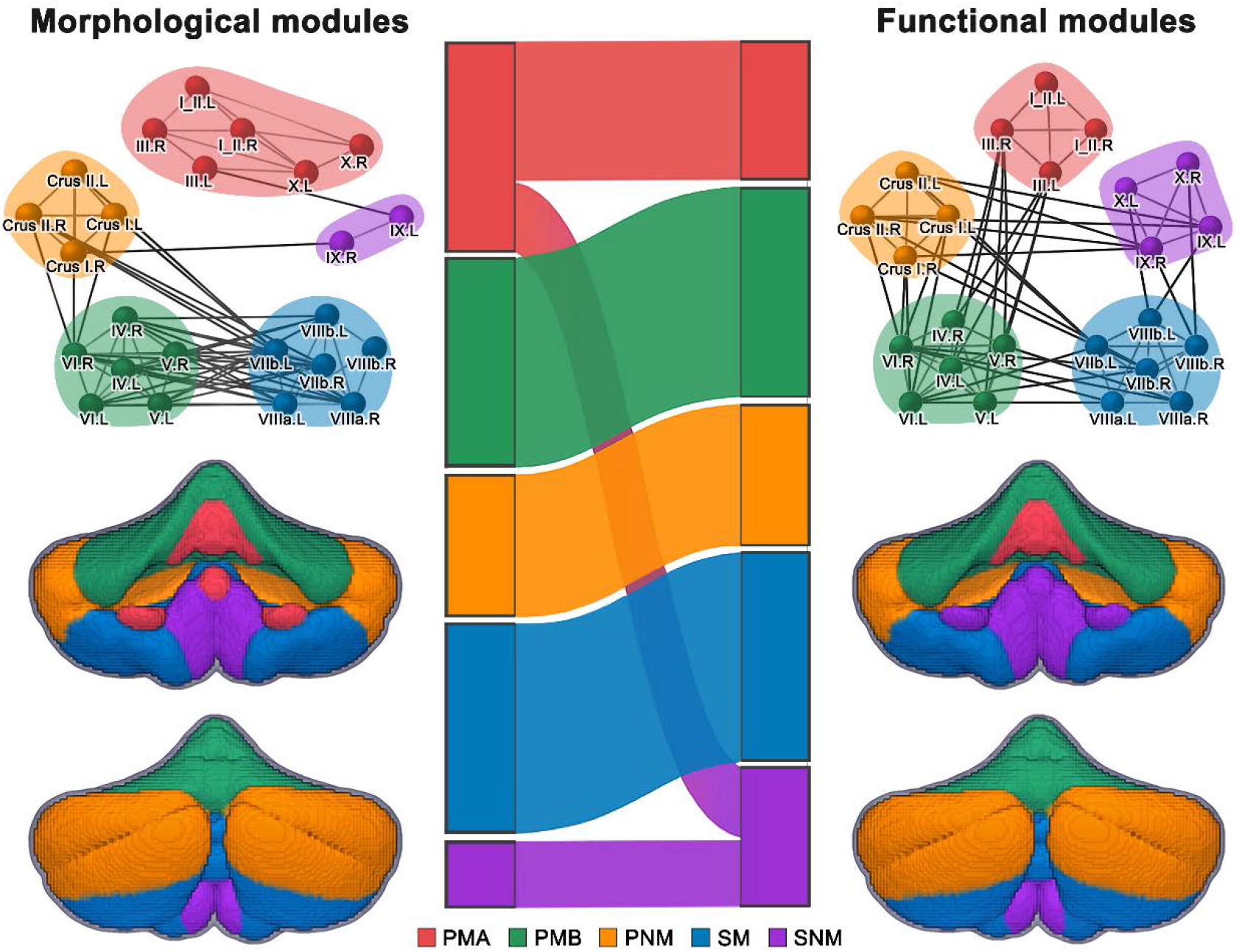
Modular architecture of the cerebellum. Five cerebellar modules were identified by applying a multilayer community detection algorithm to the group-level multiplex network of the HCs that integrated morphological and functional connectivity within the cerebellum. Module assignments of cerebellar lobules were largely comparable between morphological and functional networks. PMA, Primary Motor A; PMB, Primary Motor B; PNM, Primary Non-Motor; SM, Secondary Motor; SNM, Secondary Non-Motor.

### Alterations in cerebellar module-based cortical thickness

Significant group effects were found on the mean cortical thickness within the cerebellar PMA, PMB, SM and SNM (*p* < 0.05, FDR corrected). Post hoc comparisons revealed that the group effects were due to common cortical thickening to the two patient groups ([MS = NMOSD] > HCs): SM and SNM), MS-specific cortical thickening (MS > [NMOSD = HCs]: PMA), or NMOSD -specific cortical atrophy (NMOSD < [MS = HCs]: PMB) (Figure 4).

**Figure 4.**
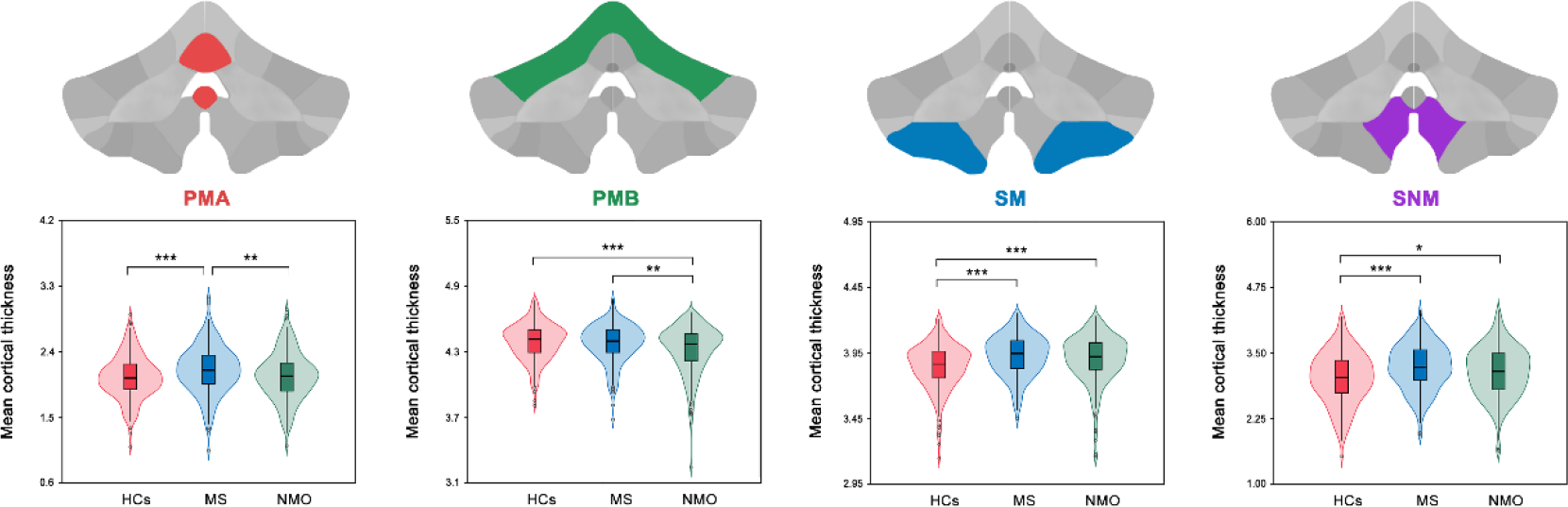
Alterations in cerebellar module-based cortical thickness. Significant group effects were found on mean cortical thickness within four cerebellar modules (PMA: MS-specific cortical thickening; PMB: NMO-specific cortical atrophy; SM and SNM: common cortical thickening to the two patient groups). PMA, Primary Motor A; PMB, Primary Motor B; SM, Secondary Motor; SNM, Secondary Non-Motor; *, *p* < 0.05; **, *p* < 0.01; ***, *p* < 0.001.

### Alterations in cerebellar morphological connectivity

Differences in cerebellar morphological connectivity are shown in Figure 5 (top panel).

**Figure 5.**
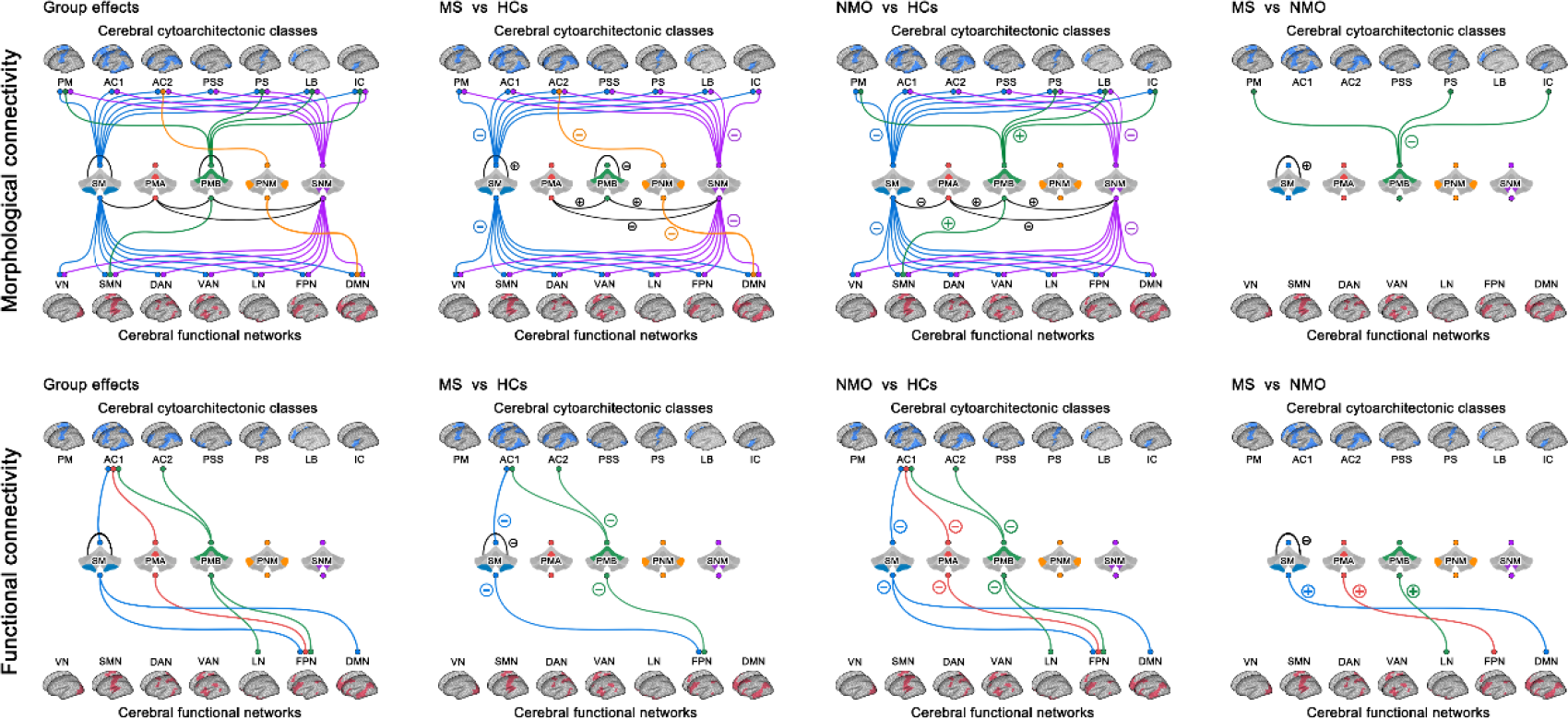
Alterations in cerebellar module-based morphological and functional connectivity. For morphological connectivity, numerous alterations of both increases and decreases were found in the two patient groups. Compared with morphological connectivity, fewer alterations were observed for functional connectivity in the two patient groups that were all characterized by disease-related decreases in particular for connectivity between cerebellar motor modules and cerebral association cortex or high-order networks. PMA, Primary Motor A; PMB, Primary Motor B; PNM, Primary Non-Motor; SM, Secondary Motor; SNM, Secondary Non-Motor; PM, primary motor cortex; AC1, association cortex; AC2, association cortex; PSS, primary/secondary sensory; PS, primary sensory cortex; LB, limbic regions; IC, insular cortex; VN, visual network; SMN, somatomotor network; DAN, dorsal attention network; VAN, ventral attention network; LN, limbic network; FPN, fronto-parietal network; DMN, default mode network.

#### Within-cerebellar morphological connectivity

Significant group effects were found on the mean morphological connectivity within two cerebellar modules (PMB and SM) and between four pairs of cerebellar modules (PMA - PMB, PMA - SM, PMA - SNM and PMB - SNM) (*p* < 0.05, FDR corrected). Post hoc comparisons revealed that the group effects were owing to MS-related decreases (MS < HCs: PMB), MS-specific increases (MS > [NMOSD = HCs]: SM), common decreases ([MS = NMOSD] < HCs: PMA - SNM) and increases ([MS = NMOSD] > HCs: PMA - PMB and SNM - PMB) to the two patient groups, or NMOSD-related decreases (NMOSD < HCs: PMA - SM).

#### Cerebello-cerebral morphological connectivity

In the context of the cerebrum^C^, significant group effects were found on the mean morphological connectivity for the cerebellar SM and SNM with all cerebral cytoarchitectonic classes, cerebellar PMB with cerebral PM, PS, LB and IC, and cerebellar PNM with cerebral AC2 (*p* < 0.05, FDR corrected). Post hoc comparisons revealed that the group effects were because of common decreases to the two patient groups ([MS = NMOSD] < HCs: SM - all cerebral cytoarchitectonic classes, and SNM - AC1, AC2, PSS, PS and LB), NMOSD-specific increases (NMOSD > [MS = HCs]: PMB - PM, PS and IC), NMOSD-related increases (NMOSD > HCs: PMB - LB), or MS-related decreases (MS < HCs: PNM - AC2, and SNM - PM and IC). When the cerebrum^F^ was used, significant group effects were found on the mean morphological connectivity for the cerebellar SM and SNM with all cerebral functional systems, cerebellar PMB with cerebral SMN, and cerebellar PNM with cerebral DMN (*p* < 0.05, FDR corrected). The group effects originated from common decreases to the two patient groups ([MS = NMOSD] < HCs: SM and SNM - all cerebral functional systems), NMOSD-related decreases (NMOSD < HCs: PMB - SMN), or MS-related decreases (MS < HCs: PNM - DMN).

### Alterations in cerebellar functional connectivity

Differences in cerebellar functional connectivity are summarized in Figure 5 (bottom panel).

#### Within-cerebellar functional connectivity

Significant group effects were found on the functional connectivity only within the cerebellar SM (*p* < 0.05, FDR corrected) due to MS-specific decreases (MS < [NMOSD = HCs]).

#### Cerebello-cerebral functional connectivity

In the context of the cerebrum^C^, significant group effects were found on the mean functional connectivity for the cerebellar PMB with cerebral AC1 and AC2, cerebellar SM with cerebral AC1, and cerebellar PMA with cerebral AC1 (*p* < 0.05, FDR corrected). Post hoc comparisons revealed that the group effects were attributable to common decreases to the two patient groups ([MS = NMOSD] < HCs: PMB - AC1 and AC2, and SM - AC1), or NMOSD-related decreases (NMOSD < HCs: PMA - AC1). When the cerebrum^F^ was used, significant group effects were found on the mean functional connectivity for the cerebellar PMB with cerebral FPN and LN, cerebellar SM with cerebral FPN and DMN, and cerebellar PMA with cerebral FPN (*p* < 0.05, FDR corrected). The group effects were driven by common decreases to the two patient groups ([MS = NMOSD] < HCs: PMB - FPN, and SM - FPN), or NMOSD-specific decreases (NMOSD < [MS = HCs]: PMA - FPN, PMB - LN, and SM - DMN).

### Relationship between imaging features and other variables in the patients

Only in the MS group, significant correlations were found characterized by positive correlations between the mean cortical thickness of the cerebellar PMA and disease duration (r = 0.193, *p* = 0.006), lesion volume (r = 0.235, *p* < 0.001) and EDSS (r = 0.207, *p* = 0.003), and between the mean cortical thickness of the cerebellar SNM and lesion volume (r = 0.230, *p* < 0.001) (Figure 6).

**Figure 6.**
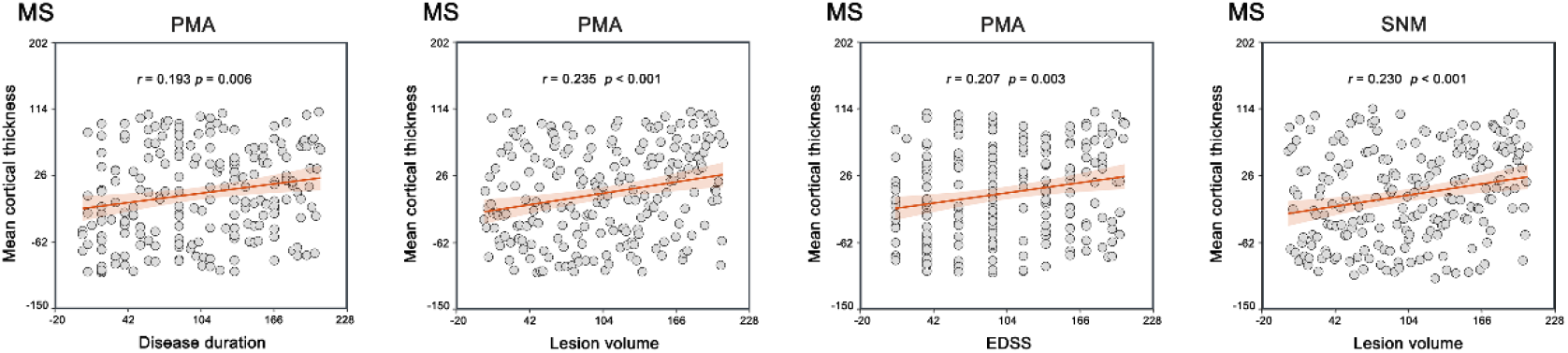
Relationships between cerebellar module-based cortical thickness and disease duration, lesion volume and EDSS in the MS patients. Among the patients, significantly positive correlations were found for mean cortical thickness within the PMA with disease duration, lesion volume and EDSS, and for mean cortical thickness within the SNM with lesion volume. MS, multiple sclerosis; PMA, Primary Motor A; SNM, Secondary Non-Motor; EDSS, expanded disability status scale.

### Classification results

The cerebellar morphological and functional connectivity could distinguish the three groups from each other with around 60% accuracy (MS vs HCs: 63.3%, *p* < 0.001; NMOSD vs HCs: 64.4%, *p* < 0.001; MS vs NMOSD: 57.9%, *p* = 0.001). As shown in Figure 7, the features contributing to the classification were mainly composed of those showing between-group differences (MS vs HCs: 83.7%; NMOSD vs HCs: 80.0%; MS vs NMOSD: 55.6%). Interestingly, morphological connectivity predominated in features contributing to the differentiation between the patients and HCs (MS vs HCs: 75.5%; NMOSD vs HCs: 64.0%) while the classification between the two diseases mainly benefited from functional connectivity (MS vs NMOSD: 88.9%). In the context of cerebellar modular architecture, the morphological connectivity contributing to the classification between the patients and HCs were mainly related to the PMB, SM and SNM (MS vs HCs: 91.9%; NMOSD vs HCs: 100%). For classifying the two patient groups, the functional connectivity contributing to the classification were all involved in the three motor-related modules (i.e., the PMA, PMB and SM; MS vs NMOSD: 100%).

**Figure 7.**
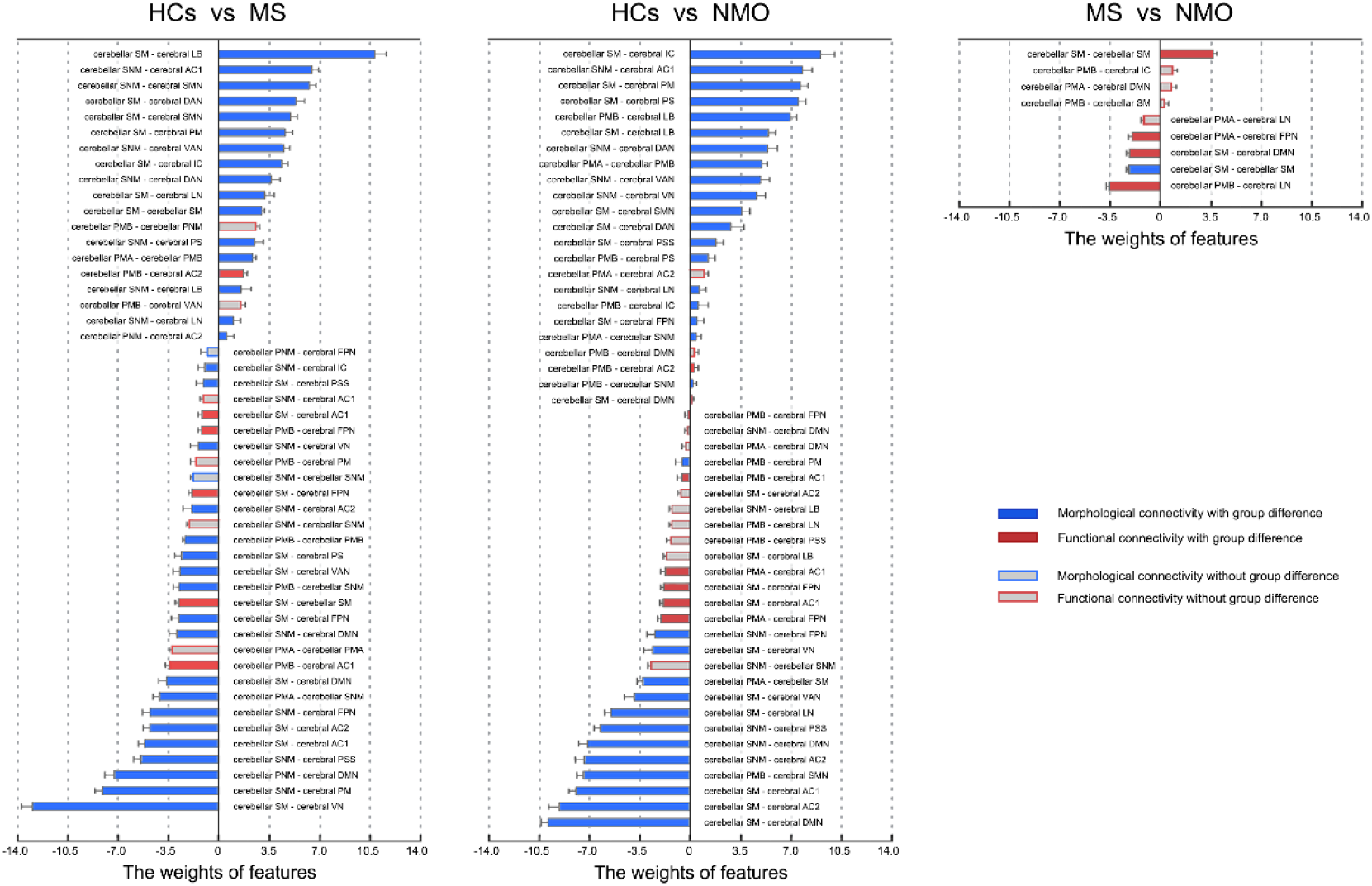
Features contributing to the classification between groups. The features contributing to the classification were mainly composed of connectivity that exhibited significant between-group differences. The classification between the patients and HCs mainly relied on morphological connectivity in particular those involving the PMB, SM and SNM, while the classification between the two diseases mainly benefited from functional connectivity that were all involved in the three motor-related modules (i.e., the PMA, PMB and SM). PMA, Primary Motor A; PMB, Primary Motor B; PNM, Primary Non-Motor; SM, Secondary Motor; SNM, Secondary Non-Motor; PM, primary motor cortex; AC1, association cortex; AC2, association cortex; PSS, primary/secondary sensory; PS, primary sensory cortex; LB, limbic regions; IC, insular cortex; VN, visual network; SMN, somatomotor network; DAN, dorsal attention network; VAN, ventral attention network; LN, limbic network; FPN, fronto-parietal network; DMN, default mode network.

## Discussion

In this study, we investigated the cerebellar morphological and functional connectivity in patients with MS and NMOSD. For cerebellar morphological connectivity, common alterations of both increases and decreases in within-cerebellar coordination and decreases in cerebello-cerebral communication were observed in the two patient groups. The MS patients showed specific increase within the cerebellar SM module while the NMOSD patients showed specific increases between cerebellar primary motor module and cerebral areas involving motor and sensory domains. Relative to morphological connectivity, fewer alterations were observed in cerebellar functional connectivity in the two patient groups that were all characterized by disease-related reductions and were mainly involved in cerebello-cerebral interactions between cerebellar motor modules and cerebral association cortex and high-order networks. The MS patients showed specific decrease within the cerebellar SM module while the NMOSD patients exhibited specific decreases between cerebellar motor modules and cerebral limbic system and default mode regions. These network-level cerebellar dysfunctions provide novel insights into shared and unique pathophysiologic mechanisms between MS and NMOSD.

### Cerebellar modular architecture

Modular organization is one of the main organizational principles of the human brain networks (Sporns and Betzel 2016). For the cerebellum, a hierarchical double-motor/triple-non-motor organization has been well established previously based on cerebellar representations of cerebral intrinsic connectivity networks (Buckner et al. 2011), cerebellar task-evoked activity (Guell, Gabrieli, and Schmahmann 2018), cerebello-cerebral functional connectivity profiles (Guell, Gabrieli, and Schmahmann 2018) and gradients of within-cerebellar functional connectivity patterns (Guell et al. 2018). In contrast to single modality in these previous studies, here we utilized a multiplex network model to merge both morphological and functional connectome information within the cerebellum for cerebellar module detection. Five modules were reliably identified with module composition largely comparable between morphological and functional cerebellar networks. Moreover, the cerebellar modular architecture well matches with the cerebellar double-motor/triple-non-motor organization, and thus provide further, in particular new anatomical, evidence for the cerebellar hierarchical organization. Specifically, the PMA and PMB correspond to the first motor representation of the cerebellum and are mainly involved in leg and foot movements ( Marek et al. 2018;; Xue et al. 2021), and hand and face movements ( Guell, Gabrieli, and Schmahmann 2018; Marek et al. 2018; Xue et al. 2021), respectively; The PNM is a mixture of the first and second non-motor representations of the cerebellum and is primarily related to working memory, executive function, language, social processing and emotion processing ( E et al. 2014; Guell, Gabrieli, and Schmahmann 2018). The SM mirrors the second motor representation of the cerebellum and is mainly engaged in motor activities with demands of task focus, visual process, attention and working memory (Brissenden et al. 2018; Guell et al. 2018); The SNM resembles the third non-motor representation of the cerebellum and is predominantly engaged in compounded domains including working memory and emotion processing, and sometimes additional visuo-spatial functions and balance maintaining (Barmack and Pettorossi 2021; Guell, Gabrieli, and Schmahmann 2018; Guell and Schmahmann 2020; Palesi et al. 2021).

### Alterations in cerebellar cortical thickness

Our main finding of cortical thickness comparisons was cortical thickening in both MS and NMOSD patients. This is in contrast to previous cerebellar studies based on gray matter volume that consistently reported decreases in MS and NMOSD (Calabrese et al. 2010; Cocozza et al. 2018; Grothe et al. 2017; Sun et al. 2019). The discrepancy is in line with the viewpoint that volumetric measures may overlook specific morphological alterations in diseases since they reflect a composite of cortical thickness, surface area and folding (Hutton et al. 2009). Interpretation of cortical thickening is not straightforward. One speculative interpretation is attributable to delays in normal physiological cleaning process caused by disease effects on pruning redundancy (Hong et al. 2016). Another possible reason, which may be more relevant to this study, is compensatory cortical reorganization to gain or maintain function (Burge et al. 2016). We found that both the MS and NMOSD patients exhibited cortical thickening in the SM and SNM modules. Given the functions of these two modules as discussed above, the common cortical thickening might reflect similar compensatory mechanisms in response to functional declines shared by the two diseases in visual scanning, motor speed, attention, learning and working memory (De Souza and Ashburn 1996; Guimarães and Sá 2012; Kawachi 2019; Oertel et al. 2019). In addition to the transdiagnostic cortical thickening, MS-specific cortical thickening was observed in the PMA module, which might be due to adaptive reorganization caused by leg restlessness, a frequent and unique sensory symptom in MS patients (Manconi et al. 2007; Schürks and Bussfeld 2013). Moreover, the PMA cortical thickness was positively correlated with EDSS scores and disease durations in the MS patients, suggesting that cortical thickening may be a biomarker to monitor clinical disability of MS. Finally, we found that the PMB module exhibited NMOSD-specific cortical thinning. Several previous case studies have reported that NMOSD patients presented choreoathetosis or pseudoathetosis, a series of involuntary and irregular movements of the face, head, hand or limbs (Boddu and Shenker 2019; Seok, Jang, and You 2018; Sugeno 2013). In addition, headaches and craniofacial pain were reported in NMOSD patients (Nielsen et al. 2018). Since cortical thinning typically reflects neuron loss and functional deficits, the thinner PMB is speculated to, at least partly, be responsible for these symptoms in NMOSD patients.

### Alterations in cerebellar morphological connectivity

Both the MS and NMOSD patients showed aberrant morphological connectivity within the cerebellum. First, the patients with MS and NMOSD exhibited common decreases in morphological connectivity between the PMA and SNM. As mentioned above, the PMA is mainly involved in leg and foot movements, while the SNM is predominately related to visuospatial function. Thus, the decreased morphological coordination may contribute to impaired visual-motor integration in MS and NMOSD (Harder et al. 2015; Julian et al. 2013; Nunan-Saah et al. 2015). Interestingly, both the PMA and SNM exhibited increased connectivity with the PMB in the two patient groups. The enhancement may reflect a common mechanism to the two diseases to compensate for pathological disruptions in the morphological connectivity between the PMA and SNM by constructing an indirect, alternative pathway. Since both the patient groups exhibited decreased morphological connectivity within the PMB (notably, the decrease did not reach significant in NMOSD after correcting for multiple comparisons), the compensatory process may occur by adaptively altering internal morphology of the PMB module in a bimodal way to enhance its morphological homogeneity with the PMA and SNM, respectively, which finally alleviates the loss of visual-motor integration in patients. Second, we found morphological connectivity increase within the SM in the MS patients. This cannot be simply interpreted as a derivative of MS-related overall cortical thickening in the SM since cortical thickening but intact morphological connectivity was observed in the NMOSD patients. Thus, the MS-specific morphological connectivity increase implies the existence of unique mechanisms that make the SM towards more similar morphology, presumably to compensate for motor-related dysfunction. The increase provides possible physiological substrate of reversibility of neurological disability in MS (Akaishi et al. 2020).

For cerebello-cerebral morphological connectivity, the two patient groups showed extensive disruptions. Specifically, the MS and NMOSD patients showed common decreases for the SM and SNM with almost all the cerebral components regardless of the cytoarchitectonic classification or functional partition. The cerebellum is closely linked with the cerebrum via complex cerebello-cerebral loops to exchange information with each other and participate collectively in various motor and non-motor activities. Thus, the widely disrupted cerebello-cerebral morphological loops may be related to poor performance on a set of motor and non-motor activities in MS and NMOSD. Notably, the disruptions were specific to the secondary rather than primary modules. Previous studies have suggested that the secondary modules are different from the primary modules in contributing to motor and non-motor activities (Guell et al. 2018; Schmahmann et al. 2019). Compared with the primary modules, the secondary modules are engaged in motor processes that require higher task focus and sometimes visual process rather than pure movements, and are devoted to processes involving multiple cognitive and affective activities rather than single non-motor domain (Guell et al. 2018; Schmahmann et al. 2019). Taking these facts into consideration, we could reasonably deduce that the common disruptions in numerous cerebello-cerebral loops to both the patient groups that were selectively involved the SM and SNM modules might account for the well described deficits in those compounded motor and non-motor activities as reflected by worse performance on BVMT, SDMT and CVLT scales in MS and NMOSD (Cao et al. 2020; Corfield and Langdon 2018; Guo et al. 2019; Meng et al. 2017).Besides the common morphological connectivity decreases, we found NMOSD-specific morphological connectivity enhancement between the cerebellar PMB module and cerebral PM, PS, IC and SMN, all of which play important roles in motor and sensory domains. The morphological connectivity increases might be seen as a compensatory response to NMOSD-specific cortical thinning in the PMB module to indemnify motor and sensory functions in patients.

Finally, we found MS-related cerebello-cerebral morphological connectivity reductions for the cerebellar PNM with the cerebral AC2 and DMN. The cerebellar PNM is constituted of the bilateral Crus I and Crus II, which are key cognitive lobules in the cerebellum and play crucial roles in working memory, attention, language, executive function and emotional self-experiences (S. H. A. Chen and Desmond 2005; E et al. 2014; Guell, Gabrieli, and Schmahmann 2018; C. Stoodley and Schmahmann 2009). Previous studies have indicated that both cerebellar Crus I and Crus II have close connection with cerebral association cortex(Sasaki et al. 1975; Xue et al. 2021), and lesions affecting the Crus I and Crus II lead to cognitive impairments by interrupting cerebellar regulation of cognitive loops with cerebral association cortex (C. J. Stoodley and Schmahmann 2010). On the other hand, the cerebellar Crus I and Crus II are also found to be major sites to interact with the cerebral DMN (Buckner et al. 2011), and damages to cerebello-cerebral DMN connections are related to cognitive impairments in MS patients (Savini et al. 2019). Accordingly, we speculate that the disrupted cerebello-cerebral loops between the cerebellar PNM and the cerebral AC2 and DMN may contribute to a variety of cognitive deficits in MS patients. Notably, no alterations in cortical thickness and within-cerebellar morphological connectivity were observed for the cerebellar PNM, suggesting a susceptibility of MS pathology to long-range connectivity of this module with the cerebrum. Using diffusion tensor images to estimate structural pathways in the cerebrum, a previous study showed that long-range connections linking remote regions were more severely disrupted in MS (Meijer et al. 2020). Expanding the previous finding, our results suggest that the vulnerability of long-range connections to MS holds for cerebellar connectivity and the susceptibility is salient for certain regions or circuits. Moreover, our findings provide important clues to a previous viewpoint that cerebellar inflammation amplifies other pathologic mechanisms in MS via long-range connections, such as retrograde neurodegeneration of cortical regions even far away from lesion sites in the cerebellum (Muthuraman et al. 2020). Deeper understanding of these findings may benefit from future research by integrating multidimensional data, animal models and computational modeling.

### Alterations in cerebellar functional connectivity

Compared with morphological connectivity, much less functional connectivity was detected to show alterations in the patients and the alterations exhibited an aggregated pattern mainly involving cerebellar motor modules and cerebral association cortex or high-order networks. Moreover, all the functional connectivity alterations were characterized by decreases in the patients. Studies related to MS pathology have suggested that moderate structural damage to the brain can be compensated by adaptive functional connectome reconfiguration, but once the structural damage exceeds a certain upper limit, the functional connectome collapses and clinical symptoms and cognitive impairments ensue (Fleischer, Radetz, et al., 2019; Schoonheim, Meijer, & Geurts, 2015). According to this theory, the observed functional connectivity disruptions may reflect functional maladaptation or failure of adaptive reorganization due to severe neurodegeneration and structural damage in the patients. Specifically, for within-cerebellar functional connectivity, MS-specific decreases were observed in the SM. According to the functions of the SM as discussed above, the decreases may be responsible for degeneration of complex motor functions and clinical disability in patients with MS.

With regard to cerebello-cerebral functional connectivity, both the patient groups showed decreases for the cerebellar PMB and SM with the cerebral association cortex and FPN. There are several lines of evidence indicating that the cerebellum is closely related with both the cerebral association cortex and the FPN. From the perspective of phylogenetic evolution, the newest parts of the cerebellum (e.g., the posterior lobe that includes the SM) develops specifically in parallel with cerebral association cortex rather than the cerebral cortex as a whole (Leiner, Leiner, and Dow 1986), facilitating the coordination between the cerebellum and cerebral association areas to exchange highly-processed multisensory information and cooperatively participate in high-level functions (Leiner, Leiner, and Dow 1991). On the other hand, from the viewpoint of functional network organization, the FPN is overrepresented in the cerebellum compared with the cerebral cortex (Marek et al. 2018), which plays a major role in task set initiation and task switching, modulating the integration of other association and motor networks (Dosenbach et al. 2007; Marek and Dosenbach 2018). Moreover, topography of the FPN in the cerebellum overlapped the cerebellar PMB and SM. Together, the disrupted functional connectivity suggests failed or weakened information exchange between the cerebellar motor modules and the cerebral association cortex as well as FPN in supporting motor, multisensory and high-level processes, which may further lead to poor performance of MS and NMOSD patients in activities involving in motor and cognition.

In addition to the common decreases, the NMOSD patients exhibited additional functional connectivity decrease for the cerebellar PMA with the cerebral association cortex and FPN, cerebellar PMB module with cerebral LN, and cerebellar SM with cerebral DMN. These findings suggest more serious disruptions in cerebello-cerebral functional integration in NMOSD, which may be related unique pathology of the disease and/or specific cognitive dysfunction of patients.

### Classification results

Our classification results indicated that cerebellar connectivity had the potential to help distinguish the patients from controls and the two diseases from each other although the accuracies were relatively low. Clinical heterogeneity of the patients, a critical concern for retrospective large-scale multisite studies, may be a main reason for the low accuracies. In addition, the accuracies can be further improved for future studies by using more sophisticated deep learning algorithms, such as convolutional neural network. Interestingly, features contributing to the classification depended on the connectivity style. This finding suggests unique susceptibility of different types of connectivity to clinical research, possibly due to their poor cross-modal correspondences as reported previously (Reid et al. 2016). Moreover, certain cerebellar modules dominated the connectivity features in the classification. Accordingly, future studies can improve the classification accuracies by exclusively focusing on a specific set of regions or connections that are closely related to the diseases.

### Limitations

This study had several limitations. First, only a subset of participants in certain sites completed a few neuropsychological assessments. This might explain why no correlations were found for the cerebellar network alterations with neuropsychological assessment. Second, there was a lack of standardized imaging protocols across different sites due to the retrospective design of this multicenter study. Although we utilized the Combat harmonization approach to mitigate site effects, it is still not clear about the extent to which our findings are contaminated by residual site effects. Third, MS and NMOSD are heterogeneous diseases both of which can be divided into different phenotypes based on clinical evolution. Future studies are thus required to explore subtype-specific cerebellar network alterations in MS and NMOSD. Finally, from a methodological viewpoint, white matter fiber tractography should be adopted in future studies which may provide additional insights beyond morphological and functional connectivity into cerebellar network dysfunctions in MS and NMOSD.

## Acknowledgements

This work was supported by the National Natural Science Foundation of China (No. 81922036), Key-Area Research and Development Program of Guangdong Province (No. 2019B030335001) and Key Realm R&D Program of Guangzhou (No. 202007030005).

