## Supplementary material for "A Multisite and Multimodal Comparative Study of Cerebellar Connectome Between Multiple Sclerosis and Neuromyelitis Optica Spectrum Disorders": Table S1

**Table S1**. Key imaging parameters in each center

| **Center** | **Scanner** | **FA (degree)** | **TR/TE (ms)** | **Voxel size (mm^3^)** | **Matrix size** | **# of Volume** |
| --- | --- | --- | --- | --- | --- | --- |
| **Functional images** | | | | | | |
| CQ | GE Discovery MR750 | 90 | 2000/40 | 3.75×3.75×4 | 64×64×33 | 240 |
| HS | GE Discovery MR750 | 90 | 2000/30 | 3.75×3.75×4 | 64×64×35 | 210 |
| JL | Siemens Skyra | 90 | 2500/30 | 3×3×3 | 70×70×43 | 200 |
| NC | Siemens Skyra | 90 | 2000/22 | 3.5×3.5×4 | 64×64×33 | 240 |
| TJ | GE Discovery MR750 | 90 | 2000/45 | 3.5×3.5×4.5 | 64×64×32 | 180 |
| XW | Siemens TrioTim | 90 | 2000/30 | 3.5×3.5×4 | 64×64×32 | 180 |
| TT | Philips Ingenia CX | 78 | 2000/30 | 3×3×4 | 80×80×40 | 180 |
| **Structural images** | | | | | | |
| CQ | GE Discovery MR750 | 12 | 8.3/3.3 | 0.5×0.5×1 | 512×512×186 |  |
| HS | GE Discovery MR750 | 12 | 8.2/3.2 | 1×1×1 | 256×256×196 |  |
| JL | Siemens Skyra | 8 | 2300/2.3 | 1×1×1 | 256×256×192 |  |
| NC | Siemens Skyra | 9 | 1900/2.3 | 1×1×1 | 256×256×176 |  |
| TJ | GE Discovery MR750 | 12 | 8.2/3.2 | 1×1×1 | 256×256×188 |  |
| XW | Siemens TrioTim | 9 | 1600/2.1 | 1×1×1 | 256×224×176 |  |
| TT | Philips Ingenia CX | 8 | 7/3 | 1×1×1 | 256×256×196 |  |

CQ, The First Affiliated Hospital of Chongqing Medical University, Chongqing; HS, Huashan Hospital Affiliated to Fudan University, Shanghai; JL, China-Japan union hospital of Jilin university, Changchun, Jinlin Province; NC, The first affiliation hospital of Nan Chang university, Nanchang, Jiangxi Province; TJ, Tianjin Medical University General Hospital, Tianjin; XW, Beijing Xuanwu hospital, Capital Medical University, Beijing; TT, Beijing Tiantan hospital, Capital Medical University, Beijing; FA, flip angle; TR, repetition time; TE, echo time.
