## Supplementary material for "A Multisite and Multimodal Comparative Study of Cerebellar Connectome Between Multiple Sclerosis and Neuromyelitis Optica Spectrum Disorders": Table S2

**Table S2.** MRI-based measures with significant group differences

|  | HCs  (n = 228) | MS  (n = 208) | NMO  (n = 200) | *p*-value |
| --- | --- | --- | --- | --- |
| **Cortical thickness** | | | | |
| PMA | 1.920 (0.348) | 2.022 (0.397) | 1.912 (0.402) | 0.002^a,b^ |
| PMB | 4.170 (0.207) | 4.143 (0.204) | 4.098 (0.248) | <0.001^a,c^ |
| SM | 3.963 (0.258) | 4.013 (0.254) | 3.997 (0.242) | <0.001^b,c^ |
| SNM | 3.103 (0.604) | 3.261 (0.540) | 3.181 (0.607) | <0.001^b,c^ |
| **Morphological connectivity** | | | | |
| Within-cerebellar | | | | |
| PMB - PMB | 0.699 (0.034) | 0.696 (0.033) | 0.695 (0.032) | 0.008^b^ |
| SM - SM | 0.678 (0.046) | 0.689 (0.045) | 0.681 (0.041) | 0.006^a,b^ |
| PMA - PMB | 0.463 (0.048) | 0.475 (0.041) | 0.478 (0.049) | 0.003^b,c^ |
| PMA - SM | 0.494 (0.060) | 0.490 (0.063) | 0.477 (0.074) | 0.004^c^ |
| PMA - SNM | 0.683 (0.110) | 0.671 (0.099) | 0.671 (0.108) | <0.001^b,c^ |
| PMA - PMB | 0.659 (0.133) | 0.691 (0.127) | 0.686 (0.136) | <0.001^b,c^ |
| Cerebello-cerebral^C^  SNM  52.0 (10.8)  47.5 (15.0)  47.0 (14.5)  <0.001^b,c^ | | | | |
| PMB - PM | 0.506 (0.046) | 0.512 (0.048) | 0.518 (0.063) | 0.007^a,c^ |
| PMB - PS | 0.472 (0.042) | 0.476 (0.039) | 0.483 (0.055) | 0.016^a,c^ |
| PMB - LB | 0.504 (0.042) | 0.507 (0.046) | 0.516 (0.052) | 0.004^c^ |
| PMB - IC | 0.497 (0.048) | 0.502 (0.047) | 0.511 (0.057) | <0.001^a,c^ |
| PNM - AC2 | 0.609 (0.068) | 0.600 (0.088) | 0.610 (0.077) | 0.025^b^ |
| SM - PM | 0.527 (0.064) | 0.519 (0.056) | 0.519 (0.046) | 0.002^b,c^ |
| SM - AC1 | 0.533 (0.060) | 0.519 (0.056) | 0.518 (0.050) | <0.001^b,c^ |
| SM - AC2 | 0.526 (0.062) | 0.512 (0.061) | 0.506 (0.054) | <0.001^b,c^ |
| SM - PSS | 0.514 (0.063) | 0.500 (0.057) | 0.497 (0.055) | <0.001^b,c^ |
| SM - PS | 0.496 (0.061) | 0.487 (0.055) | 0.486 (0.049) | <0.001^b,c^ |
| SM - LB | 0.520 (0.057) | 0.514 (0.063) | 0.513 (0.052) | 0.005^b,c^ |
| SM - IC | 0.520 (0.061) | 0.512 (0.064) | 0.512 (0.054) | 0.007^b,c^ |
| SNM - PM | 0.678 (0.075) | 0.665 (0.093) | 0.667 (0.090) | 0.008^b^ |
| SNM - AC1 | 0.680 (0.070) | 0.666 (0.087) | 0.669 (0.083) | 0.002^b,c^ |
| SNM - AC2 | 0.681 (0.079) | 0.662 (0.094) | 0.663 (0.088) | <0.001^b,c^ |
| SNM - PSS | 0.678 (0.078) | 0.655 (0.094) | 0.659 (0.091) | <0.001^b,c^ |
| SNM - PS | 0.662 (0.088) | 0.639 (0.097) | 0.639 (0.093) | 0.002^b,c^ |
| SNM - LB | 0.680 (0.078) | 0.663 (0.088) | 0.666 (0.081) | 0.004^b,c^ |
| SNM - IC | 0.674 (0.076) | 0.658 (0.088) | 0.663 (0.079) | 0.007^b^ |
| Cerebello-cerebral^F^  SNM  52.0 (10.8)  47.5 (15.0)  47.0 (14.5)  <0.001^b,c^ | | | | |
| PMB - SMN | 0.493 (0.045) | 0.497 (0.044) | 0.504 (0.059) | 0..021^c^ |
| PNM - DMN | 0.623 (0.063) | 0.615 (0.081) | 0.622 (0.069) | 0.022^b^ |
| SM - VN | 0.512 (0.053) | 0.499 (0.059) | 0.498 (0.049) | <0.001^b,c^ |
| SM - SMN | 0.515 (0.059) | 0.505 (0.057) | 0.504 (0.046) | <0.001^b,c^ |
| SM - DAN | 0.515 (0.062) | 0.506 (0.058) | 0.501 (0.054) | <0.001^b,c^ |
| SM - VAN | 0.531 (0.064) | 0.518 (0.057) | 0.513 (0.053) | <0.001^b,c^ |
| SM - LN | 0.528 (0.062) | 0.515 (0.056) | 0.514 (0.059) | <0.001^b,c^ |
| SM - FPN | 0.527 (0.062) | 0.512 (0.056) | 0.509 (0.053) | <0.001^b,c^ |
| SM - DMN | 0.539 (0.058) | 0.523 (0.057) | 0.521 (0.046) | <0.001^b,c^ |
| SNM - VN | 0.671 (0.077) | 0.652 (0.093) | 0.655 (0.088) | <0.001^b,c^ |
| SNM - SMN | 0.672 (0.082) | 0.660 (0.093) | 0.658 (0.089) | 0.006^b,c^ |
| SNM - DAN | 0.672 (0.082) | 0.658 (0.093) | 0.656 (0.090) | 0.002^b,c^ |
| SNM - VAN | 0.679 (0.072) | 0.664 (0.091) | 0.669 (0.083) | 0.002^b,c^ |
| SNM - LN | 0.686 (0.071) | 0.657 (0.089) | 0.662 (0.085) | <0.001^b,c^ |
| SNM - FPN | 0.686 (0.078) | 0.666 (0.095) | 0.667 (0.085) | <0.001^b,c^ |
| SNM - DMN | 0.683 (0.071) | 0.668 (0.083) | 0.668 (0.082) | 0.001^b,c^ |
| **Functional connectivity**  51.5 (14.0)  40.0 (17.0)  40.0 (18.0)  <0.001^b,c^ | | | | |
| Within-cerebellar  28.0 (5.8)  27.0 (11.8)  22.0 (12.0)  0.003^c^ | | | | |
| SM - SM | 0.394 (0.154) | 0.375 (0.155) | 0.419 (0.169) | <0.001^a,b^ |
| Cerebello-cerebral^C^  28.0 (5.8)  27.0 (11.8)  22.0 (12.0)  0.003^c^ | | | | |
| PMA - AC1 | -0.004 (0.077) | -0.009 (0.066) | -0.023 (0.080) | 0.003^c^ |
| PMB - AC1 | -0.008 (0.071) | -0.021 (0.055) | -0.023 (0.066) | <0.001^b,c^ |
| PMB - AC2 | -0.032 (0.087) | -0.049 (0.078) | -0.050 (0.089) | 0.003^b,c^ |
| SM - AC1 | 0.000 (0.060) | -0.014 (0.063) | -0.014 (0.056) | 0.002^b,c^ |
| Cerebello-cerebral^F^  28.0 (5.8)  27.0 (11.8)  22.0 (12.0)  0.003^c^ | | | | |
| PMA - FPN | 0.002 (0.124) | -0.001 (0.119) | -0.033 (0.130) | 0.001^a,c^ |
| PMB - LN | -0.016 (0.104) | -0.004 (0.105) | -0.044 (0.121) | 0.001^a,c^ |
| PMB - FPN | -0.040 (0.139) | -0.065 (0.114) | -0.073 (0.131) | <0.001^b,c^ |
| SM - FPN | 0.021 (0.122) | -0.006 (0.118) | -0.013 (0.098) | <0.001^b,c^ |
| SM - DMN | -0.083 (0.110) | -0.082 (0.101) | -0.106 (0.132) | <0.001^a,c^ |

Data are represented as median (interquartile range). HCs, healthy controls; MS, multiple sclerosis; NMO, neuromyelitis optica; PMA, Primary Motor A; PMB, Primary Motor B; PNM, Primary Non-Motor; SM, Secondary Motor; SNM, Secondary Non-Motor; PM, primary motor cortex; AC1, association cortex; AC2, association cortex; PSS, primary/secondary sensory; PS, primary sensory cortex; LB, limbic regions; IC, insular cortex; VN, visual network; SMN, somatomotor network; DAN, dorsal attention network; VAN, ventral attention network; LN, limbic network; FPN, fronto-parietal network; DMN, default mode network.

^a^Significant differences between the two patient groups.

^b^Significant differences between the MS patients and HCs.

^c^Significant differences between the NMO patients and HCs.
